# Extensive functional repertoire underpins complex behaviours: insights from Parkinson’s disease

**DOI:** 10.1101/823849

**Authors:** Pierpaolo Sorrentino, Rosaria Rucco, Fabio Baselice, Rosa De Micco, Alessandro Tessitore, Arjan Hillebrand, Laura Mandolesi, Michael Breakspear, Leonardo L. Gollo, Giuseppe Sorrentino

## Abstract

Rapid reconfigurations of brain activity support efficient neuronal communication and flexible behaviour. Suboptimal brain dynamics impair this adaptability, possibly leading to functional deficiencies. We hypothesize that impaired flexibility in brain activity can lead to motor and cognitive symptoms of Parkinson’s disease (PD). To test this hypothesis, we studied the ‘functional repertoire’ – the number of distinct configurations of neural activity – using source-reconstructed magnetoencephalography in PD patients and controls. We found stereotyped brain dynamics and reduced flexibility in PD. The intensity of this reduction was proportional to symptoms severity, which can be explained by beta-band hyper-synchronization. Moreover, the basal ganglia were prominently involved in the abnormal patterns of brain activity. Our findings support the hypotheses that: symptoms in PD reflect impaired brain flexibility, this impairment preferentially involves the basal ganglia, and beta-band hypersynchronization is associated with reduced brain flexibility. These findings highlight the importance of extensive functional repertoires for behaviour and motor.

## 1. Introduction

Brain functioning requires efficient reconfiguration of patterns of activity. This flexibility is essential for the coordinated engagement of brain regions, which underlies complex behaviours. Moreover, dynamic patterns of activity reflect the coordinated engagement of different systems within the brain ^1^. Accordingly, a large number of distinct patterns of activity (“functional repertoire”) indicates flexible dynamics. Growing evidence also indicates that the maximum number of spatio-temporal patterns of activity occurs at *criticality* and deviations from this state incur in a reduction of the number of patterns observed ^2^. Criticality is a highly variable, adaptive and flexible dynamical regime ^3^. It optimizes the capability of storing information ^4^, the efficient transmission of information across the brain ^3^, the response to internal fluctuations ^5^ and the detection of external stimuli ^6^; for review see ^7^.

The study of temporally resolved patterns of activation has recently drawn significant interest ^8^. Furthermore, it has been shown that they may reveal cognitive and behavioural functions ^9^. In fact, a number of behavioural functions (both motor and cognitive) require flexible dynamics, and brain pathologies might shift the brain to another state where reconfigurations are not efficient. Hence, the behavioural outcomes that rely on efficient reconfigurations may be impaired, which may result in specific symptoms. Notably, accounting for temporal variability of brain activity has improved the diagnostic classification of neurodegenerative diseases, indicating the value of time-resolved brain-network properties ^10^.

Parkinson’s disease (PD) is a severe and disabling neurodegenerative disorder. It is characterized by reduced amplitude of movements, and slowing of cognitive processes, imposing major individual and social burden ^11^. The main histopathological finding in PD is a severe nigrostriatal dopamine depletion ^12^. PD has traditionally been regarded predominantly as a motor disease. However, recent clinical and neuroimaging findings have questioned these views ^13^. For example, the first symptoms to appear are often not motor, and the clinical phenotype clearly goes well beyond the motor impairment, with other domains, such as executive functioning, involved ^14^. Structural Magnetic Resonance Imaging (MRI) has also confirmed that the regions involved in PD are more widespread than previously thought ^15^. Although functional magnetic resonance (fMRI) studies have identified impairment in the corticostriatal network in PD patients, the observed changes in functional connectivity extend into many other brain systems ^16^. Similarly, magnetoencephalography (MEG) studies have also reported widespread alterations in multiple functional connections in PD ^17^.

The brain can be represented as a network, with, at the macro-scale, brain areas as ‘nodes’ and structural or functional relationships between the brain areas as ‘edges’ ^18^, following which the topology of the network can be characterised using graph theory. A rapidly growing body of research now uses graph theory to characterize large-scale patterns of brain activation in health ^19^ and disease ^20^. Graph theory has highlighted the multifaceted nature of large-scale interactions in PD, with studies reporting more-connected networks ^21,22^, less-connected networks ^23,24^, and a combination of both ^25,26^. However, a large body of research in this area did not explore the structure of dynamic fluctuations in brain activity or connectivity between regions. Recent studies, specifically addressing dynamical changes in PD, demonstrated that patients dwelled in a hyperconnected state more frequently compared to matched controls ^27^, and showed greater network level integration in the medication-OFF compared to the ON state ^28^. These findings suggest that impaired flexibility might be deleterious and potentially also related to clinical symptomatology.

One candidate mechanism of impaired flexibility derives from alterations in excitation and synchronizability ^29,30^. Optimal flexibility requires a moderate amount of synchronizability ^31^. In contrast, hyper-synchronization can restrict the variability of brain states, entraining the dynamics onto a limited number of patterns. Conversely, too little synchronization also limits the variability of brain states with insufficient integration ^30^. This framework may be specifically relevant for PD, where hyperactivation and hypersynchronization (primarily in in the beta band) are linked to clinical disability ^32^. Furthermore, administration of L-DOPA can partly revert aberrant hypersynchronization in the beta band and relieve symptoms ^33^.

Drawing on this theory, we here hypothesized that functional activity in PD would be less flexible than in matched controls. This flexibility is estimated based on the variability of the types of patters seen in avalanches. We hypothesized that the basal ganglia would play a particularly important role in determining the occurrence of a reduced number of patterns. Furthermore, we also expected greater symptom expression to be associated with a more restricted functional repertoire.

## 2. Results

To test these hypotheses, we used source-reconstructed resting state MEG data acquired from a cohort of 39 PD patients and 38 age-matched healthy controls (see Methods for details). We analysed the size of the functional repertoire (the number of different unique avalanche configurations, see Methods) and estimated the contribution of individual brain regions to this measure of spontaneous cortical flexibility. Moreover, we correlated the individual flexibility of patients to motor and cognitive outcomes. Finally, we estimated the correlation between synchronization in the beta band and the size of the functional repertoire. Data were acquired in two brief segments of eyes-closed resting-state conditions, each of 150 seconds duration. After artefact removal and segment rejection, the average duration of combined recordings was 131.05 seconds in the PD cohort and 131.46 seconds in controls. There was no significant difference in the average recording duration (Wilcoxon rank-sum test, p=0.4539, difference of the average length = 0.41 seconds) or the number of ICA components removed in each group (KS-stats, *p* = 1.0000 and *p* = .9999 for electrocardiogram (ECG) and electrooculogram (EOG) components, respectively).We first quantified the functional repertoire of these source-reconstructed resting state MEG data, namely the total number of distinct avalanche configurations within each participant’s data. A between-group contrast revealed that PD participants expressed a restricted functional repertoire, visiting a lower number of distinct patterns in comparable amounts of time (permutation test, p=0.0088, see Fig. 1A, B).

**Figure 1.**
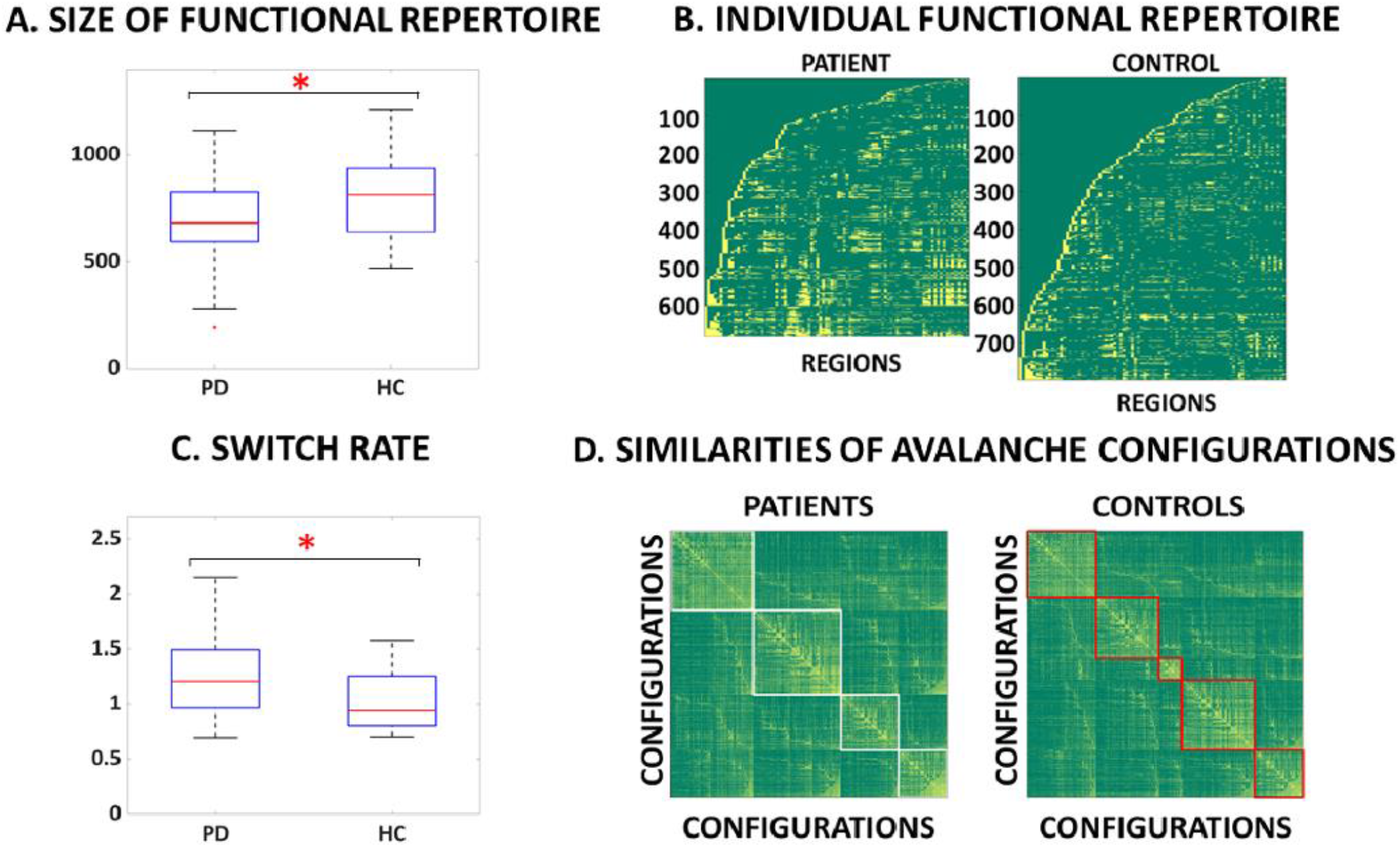
Functional repertoire computed from high spatiotemporal MEG data. **A**. The number of unique avalanche configurations in PD and healthy controls (HC). The central mark in the box indicates the median, and the edges of the box the 25th and 75th percentiles. The whiskers extend to 3 standard deviations, and the outliers are plotted individually using the ‘+’ symbol. **B**. Avalanche configurations for two representative participants (active regions in yellow, inactive regions in green). On the x-axis, the 90 AAL regions (cortex and basal ganglia). **C**. The number of switches for the two groups. **D**. Boolean similarities between avalanche configurations. Rows and colums show all avalanche configurations that are present in each cohort. The entries are the boolean similarities (i.e. yellow indicates higher similarity). The matrix-entries have been re-ordered using the Louvain modularity algorithm (as implemented in the brain connectivity toolbox), to graphically highlight the presence of clusters. The coloured lines (white and red) show the clusters in patients and controls, respectively.

The size of the functional repertoire correlated negatively with synchronization in the beta band (r = -0.4, *p* = .0070) (Fig. 2A). A negative correlation was found between the number of states visited and the clinical severity, as measured by the UPDRS-III, in the PD cohort (r(37) = -0.31, *p* = .026) (Fig. 2B).

**Figure 2.**
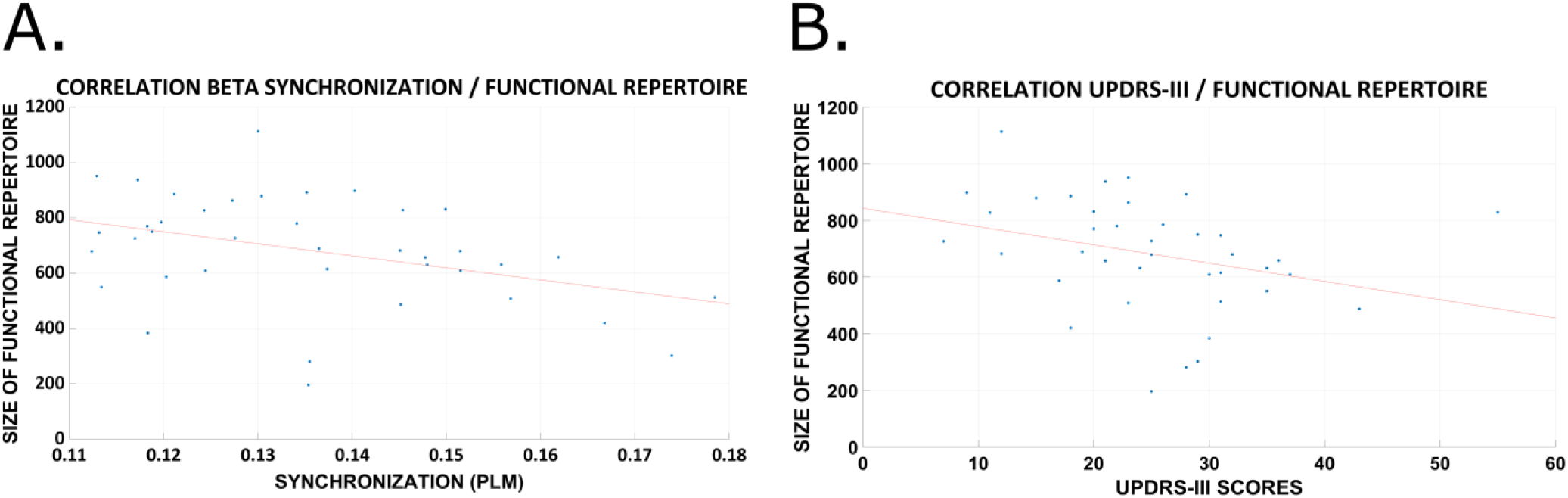
Relationship between flexibility of brain dynamics, clinical disability and global synchronization. **A**. Correlation between global synchronization, estimated using the phase linearity measurement (PLM) in the beta band, and the size of the functional repertoire. **B**. Correlation between motor impairment, estimated using the Unified Parkinson disease rating scale - III, and the size of the functional repertoire.

When relating the number of visited states with specific cognitive domains, evaluated by the MoCA subscores, linear positive relationships were present with the executive (r(37) = 0.34, *p* = .0352), attention (r(37) = 0.31, *p* = .049), language (r(37) = 0.32, *p* = .044), and naming domains (r(37) = 0.39, *p* = .015). We then compared the switch rate between activated and deactivated states across all areas, capturing the total number of activations regardless of their spatial configuration. Interestingly, PD patients showed a higher rate of switching (permutation test, *p* = .0047, Fig. 1C), implying that the decreased diversity of configurations in PD reflected more stereotyped activity, and not fewer activations *per se*.

We moved on to study the heterogeneity of avalanches across members of the same cohort, and compared this heterogeneity between the groups. Permutation testing showed that the Hamming distance across avalanche patterns belonging to either healthy controls or PD differed significantly (permutation testing, *p* = .0048), where the avalanches in the PD patients were more homogenous in comparison to controls (Fig. 1, D). This finding confirms that the ability to efficiently vary the activation patterns is impaired in PD patients. Finally, we estimated whether the distribution of the probability of each region to participate in the avalanche patterns differed between those patterns that were specific to a group vs patterns that were present in both groups. Using the Kolmogorov–Smirnov test, we observed that the two distributions were indeed statistically different (*p* = .012), implying a broad difference between regional involvement in avalanches in each group. Using bootstrapping to perform post-hoc analyses, we identified those regions that appeared more frequently in avalanche patterns that were unique to one group. The pallidum bilaterally (*p* = 0.03, *p* = 0.02, left and right, respectively), the left thalamus (*p* = 0.03), and the right putamen (*p* = 0.04) were more often involved in avalanches in the PD group than in the controls. Furthermore, the caudate bilaterally, the right thalamus and the left putamen showed differences that did not reach statistical significance (see Fig. 3).

**Figure 3.**
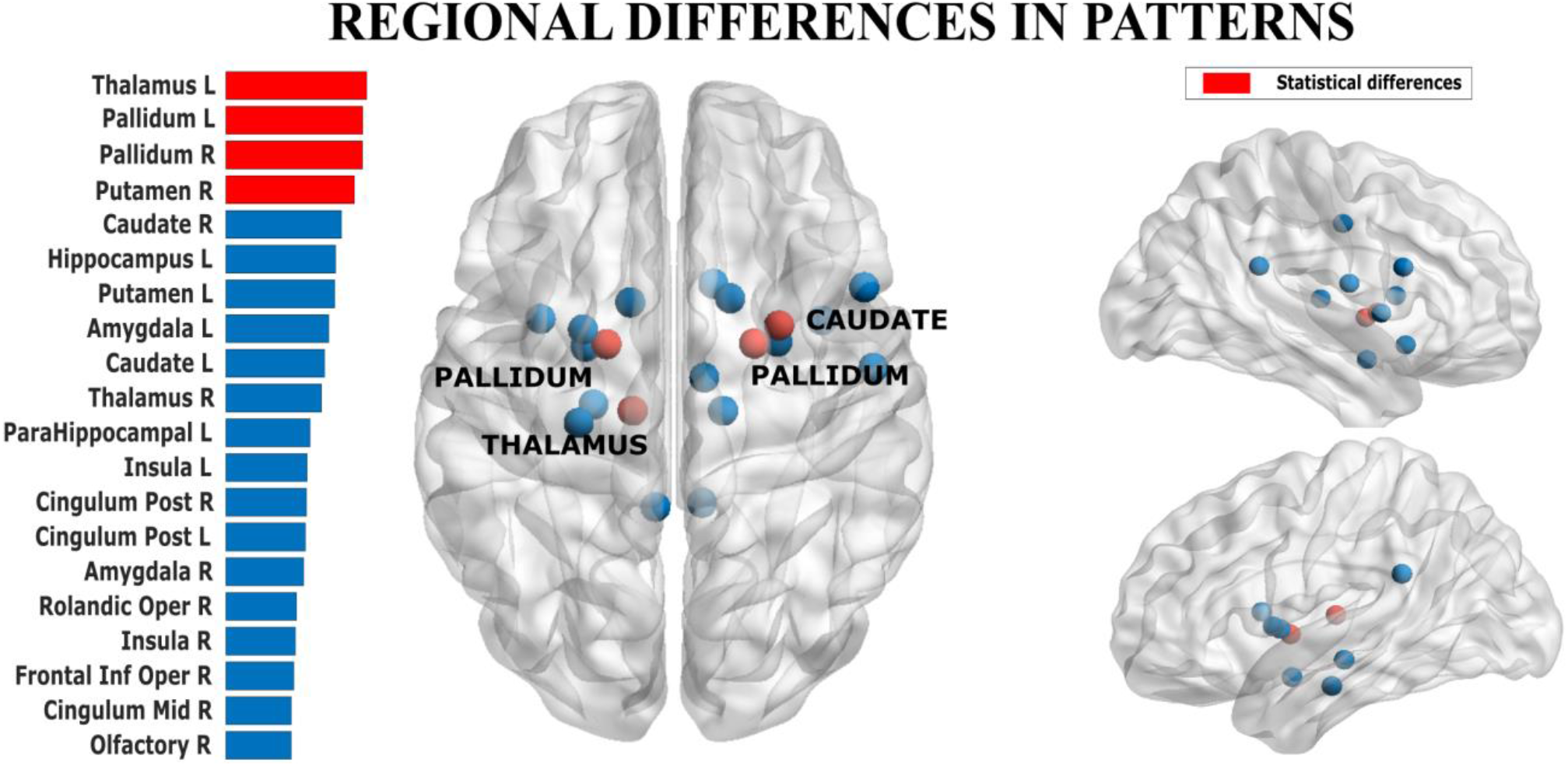
Mapping regional contribution to functional repertoire. The regions that contributed most often to avalanche patterns that were unique to one group. In the bar plot on the left, the width of the bar corresponds to the group difference in frequency with which a region contributed to the avalanche patterns that were unique to a group.

We tested the robustness of our results to specific choices of the avalanche threshold and bin length by varying these variables across a moderate range of values and repeating the analyses. Specifically, we first used different binnings, ranging from 1 to 5 (i.e. time points of the binarized time series were grouped according to the different binnings). The results remained unchanged, with patients displaying a restricted functional repertoire for all binnings explored (for the case of no binning (binning=1) and binning equal to 5, *p* = 0.011 and *p* = 0.0068, respectively; see supplementary Fig.1). Furthermore, the avalanche threshold was modified, ranging from 2.5 to 3.5. For both cases the differences between the groups were confirmed (for z=2.5 and 3.5, *p* = 0.0084 and *p* = 0.0082, respectively; see supplementary Fig. 2). We also studied the impact of including cerebellar sources, and the difference in functional repertoire remained significant (*p* = 0.0042; see supplementary Fig. 3). The analysis was also repeated for the classical frequency bands (i.e. delta, 0.4 – 4 Hz, theta 4 – 8 Hz, alpha 8 – 13 Hz, beta 13-30 Hz, gamma 30 – 48 Hz). In all the frequency bands, the difference in functional repertoire remained significant (*p* = 0.049, *p* = 0.020, *p* = 0.0065, *p* = 0.028, *p* = 0.026, from delta to gamma, respectively; supplementary Fig. 3). These results suggest that the restriction of the functional repertoire in PD is a widespread phenomenon in terms of the range of frequencies involved. Finally, when repeating the analysis using the exact same amount of data for all subjects, the group difference in functional repertoire remained significant (p<0.0001, for broadband; see supplementary Fig. 4).

## 3. Discussion

Here we investigated time-resolved flexibility of brain activity in PD, utilizing avalanches and tools from statistical physics, adapted to MEG activity. The functional repertoire was formed by distinct patterns of activation during avalanches, and the size of the functional repertoire was used as an indicator of the ease of transitions between different brain states, and hence cognitive flexibility ^7^. We found that PD patients exhibited a reduced functional repertoire compared to healthy controls. Moreover, the clinical disability was more pronounced in patients whose functional repertoire was more impoverished. Finally, we demonstrated the robustness of our results across a range of parameters and frequency bands, showing that the results did not depend on specific details of data processing.

Our results showed that PD patients display a restricted functional repertoire as compared to healthy controls. This was demonstrated by the lower number of distinct avalanche configurations. We focused on the flexibility of the brain activity, since efficient reconfiguration of active areas, avoids stereotyped, repetitive dynamics, and has been shown to be important for brain functioning ^34^. The qualitative restriction in the number of patterns of activation explored by PD patients reflects the effect of pathophysiological changes on the large-scale brain dynamics. A recent EEG study that addressed large-scale dynamics found that the duration, rather than the specific order, of functional states related with symptomatology in patients with Lewy body dementia ^35^, which is a disease that shows clinical and pathological similarities to PD ^36^. However, the same relationship could not be found in PD patients ^35^. Our results add to these findings, indicating that the functional repertoire itself is relevant in PD and related to the clinical picture. In our study, we also tested if the restricted functional repertoire was a result of a slower rate of change of state (i.e. between active or inactive) of each region. Interestingly, we found a higher switching rate in PD compared to the healthy controls, suggesting that the restriction of the functional repertoire in PD is qualitative in nature, and not due to a slowing of the regional rate of changes between active and inactive states. The length of the analyzed time series did not differ significantly between the two groups, thus this result could not be explained by the duration of the acquisitions. Importantly, our results replicated when frequency–specific signals were analysed, showing that the slowing of activity that is normally seen in PD also cannot explain the observed differences in the size of the functional repertoire.

We analyzed shared and group-specific avalanche patterns. We classified any pattern as ‘shared’ if it occurred in both groups, and as ‘group-specific’ if it occurred in either group, in order to test whether some regions are specifically important in defining pathological activity patterns. The basal ganglia belonged to group-specific patterns more often than expected by chance, suggesting that the network of regions that is dynamically connected with the basal ganglia is altered in Parkinson’s disease. This finding highlights the importance of a network perspective on brain activity, whereby the basal ganglia show their role in defining the whole-brain dynamics and functional repertoire. This evidence speaks to a vast literature on the basal ganglia as part of a distributed forebrain network with a prominent role in innate behavioral routines as well as learning (retrieving) new (previously acquired) functional repertoires ^37^. Previous evidence has shown that extreme specificity is necessary in the connections between the basal ganglia, thalamus and cortex ^38^, and that the basal ganglia play a prominent role in the fine tuning of such connections ^39^. Our results suggest that the functional role of the basal ganglia might contribute to appropriate dynamical recruitment selection of brain regions.

Importantly, we tested if the restriction of the functional repertoire related to behavioral outcomes. In fact, according to our hypothesis, we expect that impaired brain dynamics at rest would underpin reduce capability to accomplish to a variety of complex tasks. The Unified Parkinson’s disease rating scale-III (UPDRS-III) is widely used to assess PD patients, and it provides a quick account of motor symptoms severity in PD. In line with our hypothesis, an inverse relationship was evident between the restriction of the functional repertoire and the UPDRS-III. This evidence supports the interpretation of the restricted functional repertoire as pathological, and points towards the idea that an impaired regulation of the brain dynamics by the basal ganglia underlies clinical impairment in PD. In the same line of thinking, the positive correlations that were present between performance on specific cognitive domains and the size of the functional repertoire, confirms that flexible activity is important for performance on a number of behavioral tasks, not limited to the motor domain. Finally, these result highlight the importance of future longitudinal studies to test the performance of the size of the functional repertoire as a potential biomarker of disease progression.

With regard to the mechanisms underpinning the lack of flexibility, these might involve perturbations in the ability to synchronize different areas ^30^. Since PD has often been associated with excessive synchronization ^40^, we expected to find a reduction in the size of functional repertoire for patients with a more pronounced hypersynchronization in the beta band, which is classically involved in PD ^33^. Our results confirmed this predicted negative relation between the number of different patterns of activation and the overall synchronization in the beta band.

Electrophysiological studies in animal models of PD ^32^ and in patients with PD have shown that dopaminergic depletion can increase oscillatory activity in the thalamo-cortical circuitry. Furthermore, while this activity has been described in multiple frequency bands, the beta band is specifically relevant to PD ^33^. The extent of such aberrant beta synchronization has been related to clinical disability, and L-DOPA has been shown to partly revert this phenomenon and relieve symptoms ^33^. Accordingly, EEG data from PD patients showed that the lack of dopamine increases coupling strength in the basal ganglia circuitry ^40^. Within this framework, our results indicate that a hypersynchronized state is deleterious as it reduces the flexibility of brain activity. In fact, excessive synchronization reduces the variability of the behaviour of the system, hence narrowing the number of states that are readily accessible ^30,41^. Hypersynchronization and hyperconnected topology have been associated with a variety of neurological diseases ^42,43^, as well as in preclinical conditions that carry an increased risk of neurodegeneration ^44^ hence investigations of the functional repertoires in these diseases and conditions could provide answers to this question.

Some limitations of our work should be considered when interpreting our results. First, we utilised the automated anatomicl labelling (AAL) atlas when reconstructing the neuronal activity, which is well suited to MEG studies but has a somewhat coarse spatial resolution. Taking into account that this work is based on MEG signals, it is important to consider that more fine-grained atlases might go over the nominal resolution of the source-reconstructed signal, which could lead to spurious results. Interestingly, parcellations that are based on MEG resting state data contain roughly the same number of parcels as the AAL atlas ^45^. Hence, we believe that this choice of atlas yields a reasonable compromise between obtaining high spatio-temporal resolution while avoiding an overestimation of the coactivations that might be induced by atlas with a too high spatial resolution. Furthermore, we chose to focus our main analysis on 90 regions, excluding the cerebellum, as the parcellation of the cerebellum in the AAL atlas is particularly fine grained, the posterior fossa is prone to artefact, and MEG does not provide high spatial resolution. However, the role of the cerebellum is undoubtedly important in PD ^46^. For this reason, we also repeated the analysis with all 116 AAL regions, including the cerebellum. All the significant group differences were confirmed, proving the robustness of the findings. Furthermore, we chose to include the basal ganglia in the analysis, given the paramount importance of their role in PD. Nonetheless, it is worth to consider that the sensitivity of MEG, and thereby the quality of the source reconstruction, tends to decay with the depth of the analysed brain structure. Hence, the analysis involving the basal ganglia is based on a reconstructed signal that may have suboptimal resolution (as compared to the cortical data). However, it has recently been shown that magnetoencephalography can indeed detect the activity of deep brain sources ^47,48^.

In conclusion, our results show that the brain of PD patients is less flexible, and a more stereotyped brain activity impairs the ability of the brain to perform complex tasks, and is related to hypersynchronization of brain activity. Crucially, the size of the functional repertoire is proportional to the observed clinical disability. Within this framework, main symptoms of PD patients can be explained, and the known role the basal ganglia play in defining PD pathophysiology naturally emerges in the marked involvement in pathological configurations. Our findings also indicate that the mechanisms underlying impaired flexibility of brain activity relates with the observed clinical phenotype observed in PD, and to hypersynchronization. This also provides a new interpretation of PD pathophysiology in which the well-known beta hypersynchronization is associated with reduced flexibility of brain dynamics.

## 4. Materials and methods

### 4.1 Cohort description

Consecutive early PD patients, diagnosed according to the diagnostic criteria of the UK Parkinson’s Disease Society Brain Bank Diagnostic Criteria ^49^, were recruited at the Movement Disorders Unit of the First Division of Neurology at the University of Campania “Luigi Vanvitelli” (Naples, Italy). Inclusion criteria were: a) PD onset after the age of 40 years, to exclude early onset parkinsonism; b) a modified Hoehn and Yahr (H&Y) stage ≤ 2.5. Exclusion criteria were: a) dementia associated with PD according to consensus criteria ^50^; b) relevant cognitive impairment, as evidenced by age- and education-adjusted MoCA score lower than or equal to the Italian cut-off score ^51^; c) any other neurological disorder or clinically significant or unstable medical condition (Table 1). Disease severity was assessed using the Hoehn and Yahr (H&Y) stages ^52^ and the UPDRS III ^53^. Motor clinical assessment was performed in “off-state” (off-medication overnight). Levodopa equivalent daily dose (LEDD) was calculated for both dopamine agonists (LEDD-DA) and dopamine agonists + L-dopa (total LEDD) ^54^. Global cognition was assessed by means of Montreal Cognitive Assessment (MoCA) ^55^. MoCA consists of 12 subtasks exploring the following cognitive domains: (1) memory (score range 0–5), assessed by means of delayed recall of five nouns, after two verbal presentations; (2) visuospatial abilities (score range 0–4), assessed by a clock-drawing task (3 points) and by copying of a cube (1 point); (3) executive functions (score range 0–4), assessed by means of a brief version of the Trail Making B task (1 point). All participants signed informed consent. The study was approved by the Local Ethics Committee of University of Naples “L. Vanvitelli” and was conducted in accordance to the Declaration of Helsinki (Table 1).

**Table 1:** Demographic and clinical features of PD patients.

|  | HC (n=38)<br>mean±SD | PD (n=39)<br>mean±SD | p-value |
| --- | --- | --- | --- |
| Age | 62.05±9.40 | 64.23±9.36 | NS |
| Sex (M/F) | 22/16 | 23/16 | NS |
| Disease duration (months) | - | 33.0±15.4 | - |
| H&Y stage | - | 1.8±0.5 | - |
| UPDRS III | - | 25.0±9.5 | - |
| MoCA (total) | - | 22.4±3.3 | - |
| - Memory | - | 1.5±1.5 | - |
| - Visuospatial abilities | - | 2.4±1.2 | - |
| - Executive functions | - | 2.3±1.4 | - |
| - Attention, concentration and working memory | - | 4.9±1.3 | - |
| - Language | - | 4.9±1.2 | - |
| - Temporal and spatial orientation | - | 5.9±0.4 | - |
| BDI | - | 5.2±6.2 | - |
| LEDD total | - | 289.9±152.3 | - |
| LEDD DA | - | 78.3±129.1 | - |

Data are given as mean± standard deviation (SD). NS: not significant; PD: Parkinson's disease; HC: healthy controls; H&Y: Hoehn & Yahr; UPDRS: Unified Parkinson's Disease Rating Scale; MoCA: Montreal Cognitive Assessment; BDI: Beck depression inventory; LEDD: Levodopa Equivalent Daily Dose; DA: dopamine-agonist.

### 4.2 MEG acquisition

Magnetoencephalographic data were acquired in a 163-magnetometers MEG system ^56^, with 9 reference sensors, located in a magnetically shielded room (AtB Biomag UG, Ulm, Germany). The position of four position coils and four reference points (nasion, right and left pre-auricular and apex) were digitized before acquisition, using Fastrak (Polhemus®). MEG data were acquired during two eyes closed resting state segments each 2.5 minutes long. Participants were in off-state, and were requested to relax with eyes closed and not to think of anything in particular. Instructions were delivered immediately prior to each recording via an intercom. Head position was recorded at the start of each recording segment. After an anti-aliasing filter, the data were sampled at 1024 Hz. A 4^th^-order Butterworth IIR band-pass filter was then applied to remove components below 0.5 and above 48 Hz. The filter was implemented offline using MatLab scripts within the Fieldtrip toolbox 2014 ^57^. Electrocardiogram (ECG) and electrooculogram (EOG) data were also recorded.

### 4.3 MRI acquisition

MR images were acquired on a 3-T scanner equipped with an 8-channel parallel head coil (General Electric Healthcare, Milwaukee, WI, USA) either after, or a minimum of 21 days (but not more than one month) before, the MEG recording. Three-dimensional T1-weighted images (gradient-echo sequence Inversion Recovery prepared Fast Spoiled Gradient Recalled-echo, time repetition = 6988 ms, TI = 1100 ms, TE = 3.9 ms, flip angle = 10, voxel size = 1 × 1 x 1.2 mm3) were acquired.

### 4.4 Preprocessing

A principal component analysis (PCA) was performed to reduce the environmental magnetic noise ^58^. Specifically, the filter was obtained by orthogonalizing the reference signals to obtain a base, projecting the signals from the brain sensors on this noise-base, and subsequently removing these projections in order to obtain clean data ^59^. We adopted the PCA filtering implementation available within the Fieldtrip Toolbox ^57^. The noisy segments of acquisition were identified through visual inspection of the whole dataset by an experienced rater (RR). On average, 130 ± 2 channels were used. After that, Independent component analysis (ICA) ^60^ was also performed to eliminate ECG (typically 1-2 two components) and EOG (0-1 components) contributions to the MEG signals.

### 4.5 Source reconstruction

Source reconstruction of channel data was performed using a beamforming procedure implemented in the Fieldtrip toolbox ^57^. First, the subject’s fiducial points were used to co-register the MEG data to the native subject-specific MRI. Second, using a single shell volume conduction model ^61^ and an equivalent current dipole source model, a Linearly Constrained Minimum Variance (LCMV) beamformer ^62^, based on the whole pre-processed, broad-band data, was used to reconstruct time series related to the centroids of 116 regions-of-interest (ROIs), derived from the Automated Anatomical Labeling (AAL) atlas ^63,64^. Both the atlas and the MRI were aligned to the head coordinates. We focussed on the first 90 ROIs, excluding those in the cerebellum given the low reliability of the reconstructed signal in this region. For each source, we projected the time series along the dipole direction that explained most variance by means of singular value decomposition (SVD). For each subject, we visually inspected the source-space data to check that no remaining artefact was present at the source-level.

### 4.6 Analysis of dynamics

#### 4.6.1. Neuronal avalanche and avalanche configuration

To quantify spatio-temporal fluctuations of activity, we first estimated “*Neuronal Avalanches*”. To start, each of the 90 source-reconstructed signals were z-transformed (Fig. 4A). Subsequently, each time series was thresholded according to a cut-off of 3 standard deviations (i.e. *z* =3). The robustness of the results to changes in this threshold were also investigated. We defined a neuronal avalanche as a fluctuation of activity starting when at least one region becomes active (|*z*| >3) and continuing as long as any region remains active (Fig. 4B).

**Figure 4.**
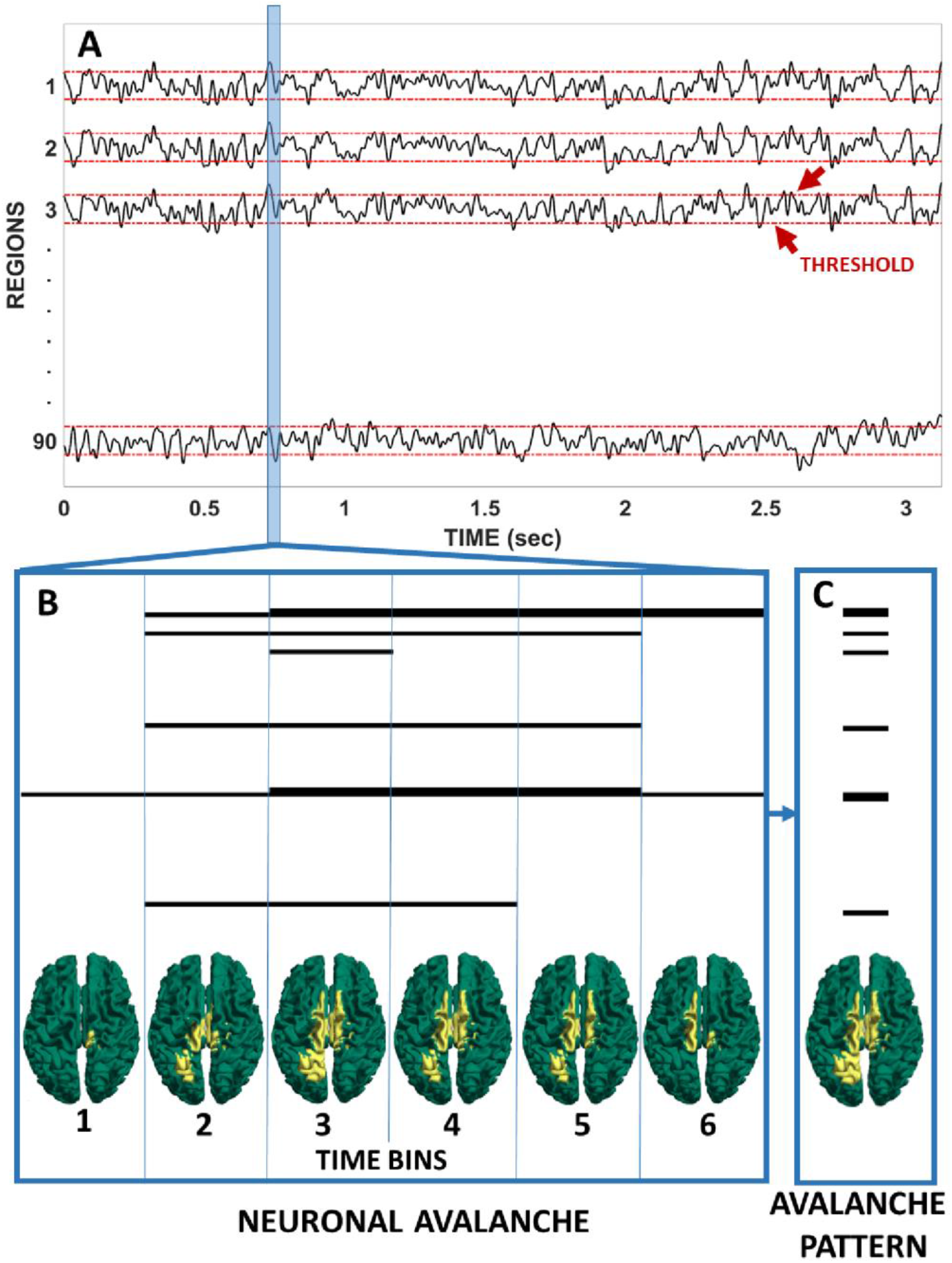
Schematic representation of an avalanche pattern. **A**. Reconstructed time series (z-scores). The red dashed lines indicate the threshold to define activation. **B**. An avalanche is a sequence of activations that starts when one or more regions are active and ends when no region is active anymore. The brains-plots for each moment (bin) of the avalanche show the active (yellow) and inactive (green) areas C. The areas that were active at some time during the avalanche together form the avalanche pattern.

These analyses require the time series to be binned. To select a suitable bin length, we computed the branching ratio ^2,65^, *σ*, as follows: for each time bin duration, for each subject, for each avalanche, the (geometrically) averaged ratio of the number of events (activations) between the subsequent time bin and that in the current time bin was calculated as,

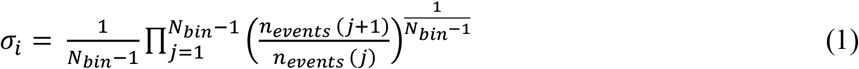

where *σ*_*i*_ is the branching parameter of the i-th avalanche in the dataset, *N*_*bin*_ is the total number of bins in the i-th avalanche, *n*_*events*_ (*j*) is the total number of events in the j-th bin. We then (geometrically) averaged the results over all avalanches ^66^,

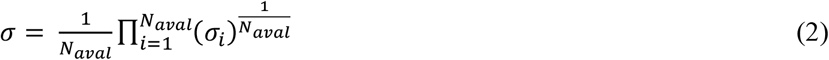

where *N*_*aval*_ is the total number of avalanche in each participant’s dataset. In branching processes, a branching ratio of *σ*=1 indicates critical processes with activity that is highly variable and nearly sustained, *σ* < 1 indicates subcritical processes in which the activity quickly dies out, and *σ* > 1 indicates supercritical processes in which the activity increases. The bin length equal to three yielded a critical process with *σ*=1, justifying the use of the term “avalanche” for these events. However, the robustness of the results to changes in this exact bin length was also investigated, and the main results were not affected (supplementary material). For each avalanche, an “*avalanche configuration*” was defined as the set of all areas that were active at some point during the avalanche (see Fig. 4C).

#### 4.6.2 Functional repertoire

For each subject, we estimated the functional repertoire, defined as the number of unique avalanche configurations that was expressed during the recording. Before comparing the functional repertoires between groups, the duration of the acquisitions was compared between the two groups, as the acquisition duration could affect the estimate of the functional repertoire. Furthermore, the analysis was also repeated with the (exact) same amount of data for each participant (67.01 seconds). To do so, we selected all participant who had at least a 1 min long acquisition (two patients had less and had to be excluded, leading to 38 controls and 37 patients). We randomly selected data segments from the whole recordings. The results from analysis of these data confirmed our initial analysis (supplementary Fig. 4).

#### 4.6.3 Switching between states

We defined a “switch” as a change of state (“on” to “off” or vice-versa) between two consecutive time – bins, occurring in any area. The switch rate (number of switches over duration), averaged over areas, was computed for each individual.

#### 4.6.4 Analysis of similarity

To estimate and compare the variability of avalanche configurations between groups, we first computed the similarities between each configuration, yielding one distance matrix per group ^2^. In each matrix, rows and columns are equal to unique avalanche patterns, while the entries are the Hamming distances (the number of regions that differ-i.e. active or inactive - in two given patterns). Statistical comparison of the similarity matrices belonging to the two groups was achieved by permutation testing (see below for details).

#### 4.6.5 Quantifying the influence of specific regions on avalanche configurations

We split the total functional repertoire in two groups: configurations that occurred in both the clinical and control participants (“shared repertoire”), and configurations that were unique to either group (“group-specific repertoire”). We then computed how often each region occurred in both shared and group-specific configurations. The Kolmogorov – Smirnov test was used to compare the two distributions, and resampling then allowed identification of those regions that appeared in group-specific repertoires more often than by chance.

### 4.7 Synchrony estimation

To estimate synchronization in the beta band, the broadband source-reconstructed data were band-pass filtered with a fourth order Butterworth filter between 13 and 30 Hz. A metric recently developed by our group ^67^, the phase linearity measurement (PLM), illustrated in Fig. 5, was then employed to estimate synchronization,

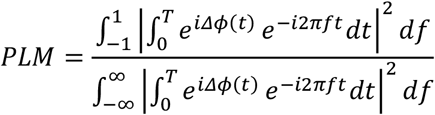

where *Δϕ*(*t*) is the interferometric phase (i.e. the phase of the signal resulting from the multiplication of one signal with the complex conjugate of another signal). The PLM was computed using the implementation available in the Fieldtrip toolbox in Matlab. In short, this metric quantifies the synchronization between two time–series using the central peak in the frequency spectrum of the interferometric signal. The more peaked the spectrum of the interferometric signal, the more the two originating signals will be synchronized. The PLM ranges between 0 and 1, is insensitive to volume conduction and grows monotonically with synchronization ^67^.

**Figure 5.**
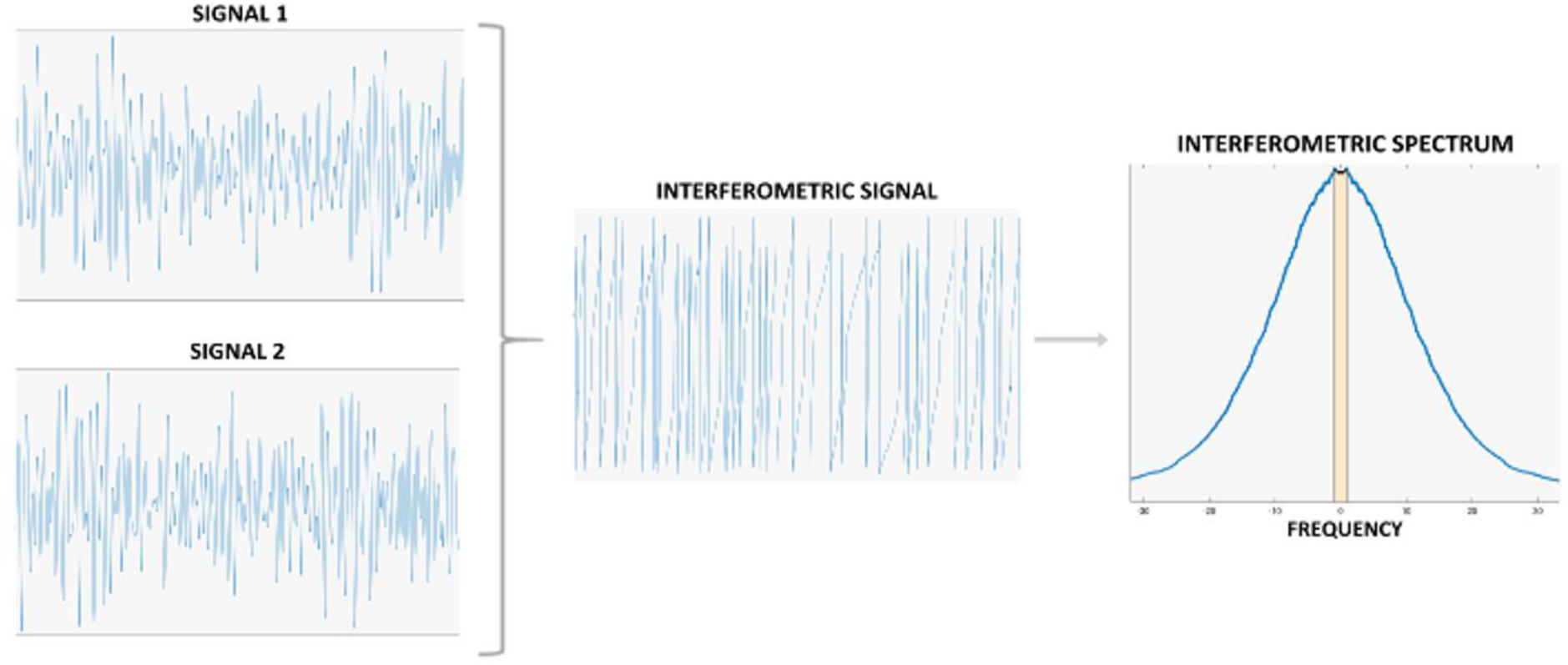
Phase linearity measurement, used to estimate synchronization. The time series of each pair of regions have been processed to obtain the interferometric signal. The power spectrum of the interferometric signal was estimated, and the central area underneath the spectrum (depicted in light brown) was compared to the total area. The higher such ratio, the more synchronized the two signals.

### 4.8 Statistical analysis

To compare age and sex between the two groups we used T-test and Chi-square test, respectively. Permutation testing or Kolmogorov-Smirnov test were performed to compare patients and controls, as appropriate. For permutation testing, the data where permuted 10000 times, and at each iteration the absolute value of the difference between the two groups was observed, building a null distribution of absolute differences. Finally, the empirical, observed difference was rank-ordered against this distribution, yielding a significance value. All statistical analyses were performed using custom scripts written in Matlab 2018a.

## Author contributions

PS and RR collected and acquired the dataset, processed the data and conceptualized the study; FB processed the data; RDM and AT collected the sample; AH, LM, MB contributed to interpreting the results and critically revised the article; LLG conceptualized and supervised the study, and GS supervised the study. All authors interpreted the results and wrote the manuscript.

## Data availability

The MEG data and the reconstructed avalanches are available upon reasonable request to the corresponding author, conditional on appropriate ethics approval at the local site.

## Supplementary Figures

**Supplementary figure 1.**
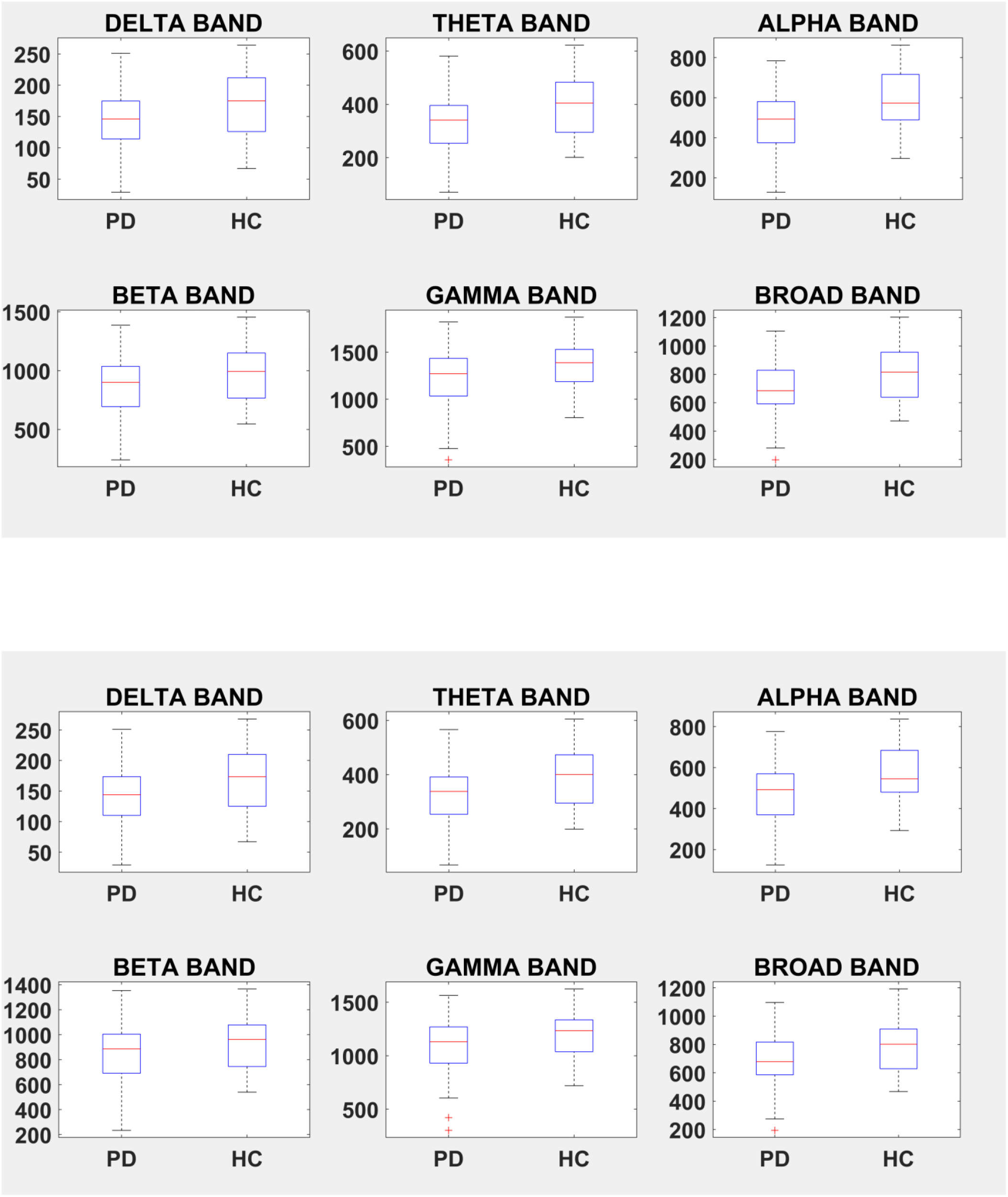
Differences in the size of functional repertoire in Parkinson patients (PD) and healthy controls (HC) with different binnings. On the top, no binning (binning = 1). On the bottom, binning = 5.

**Supplementary figure 2.**
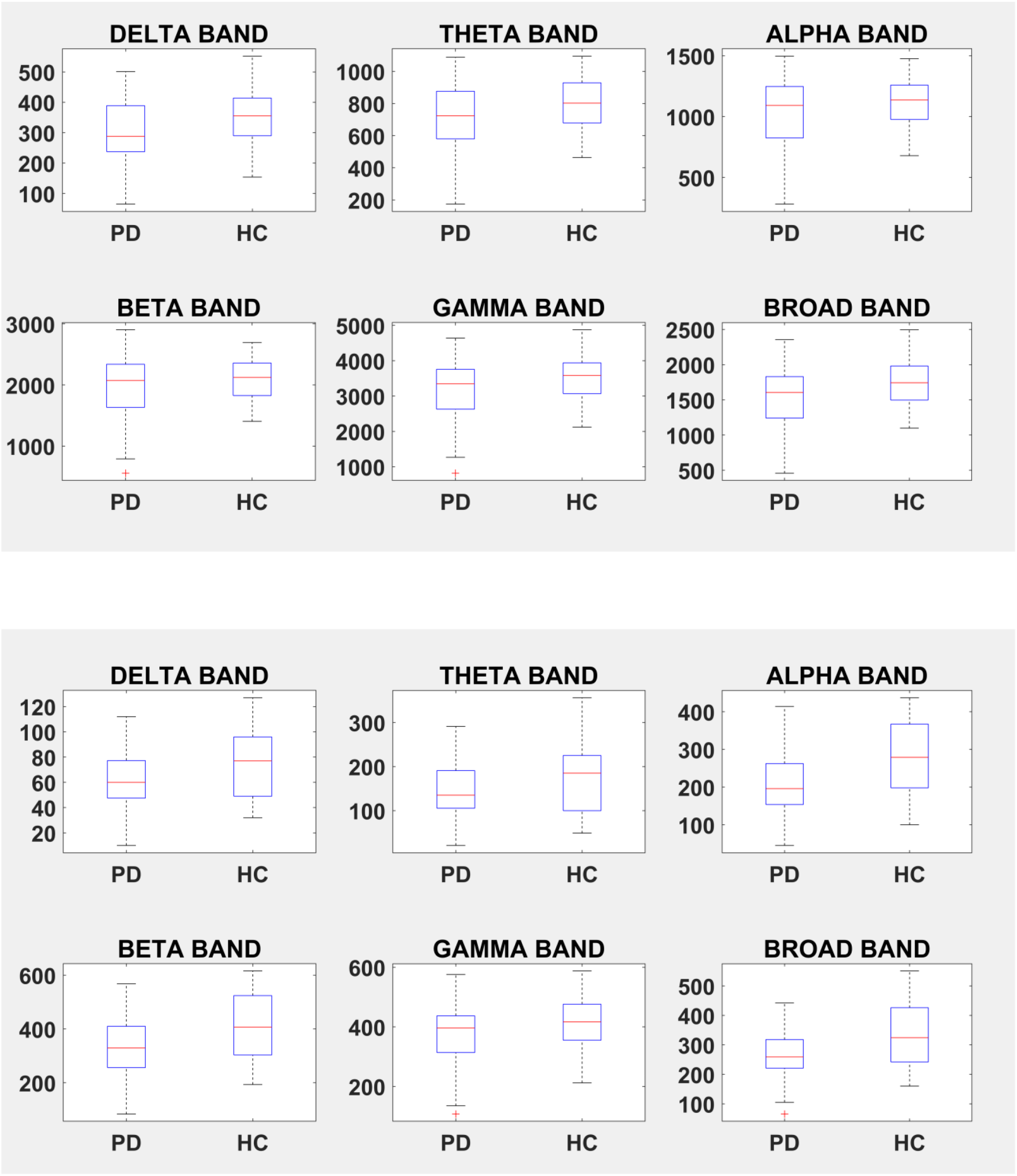
Differences in the size of the functional repertoire in Parkinson patients (PD) and healthy controls (HC) with different thresholds. On the top, threshold = 2.5. On the bottom, threshold = 3.5.

**Supplementary figure 3.**
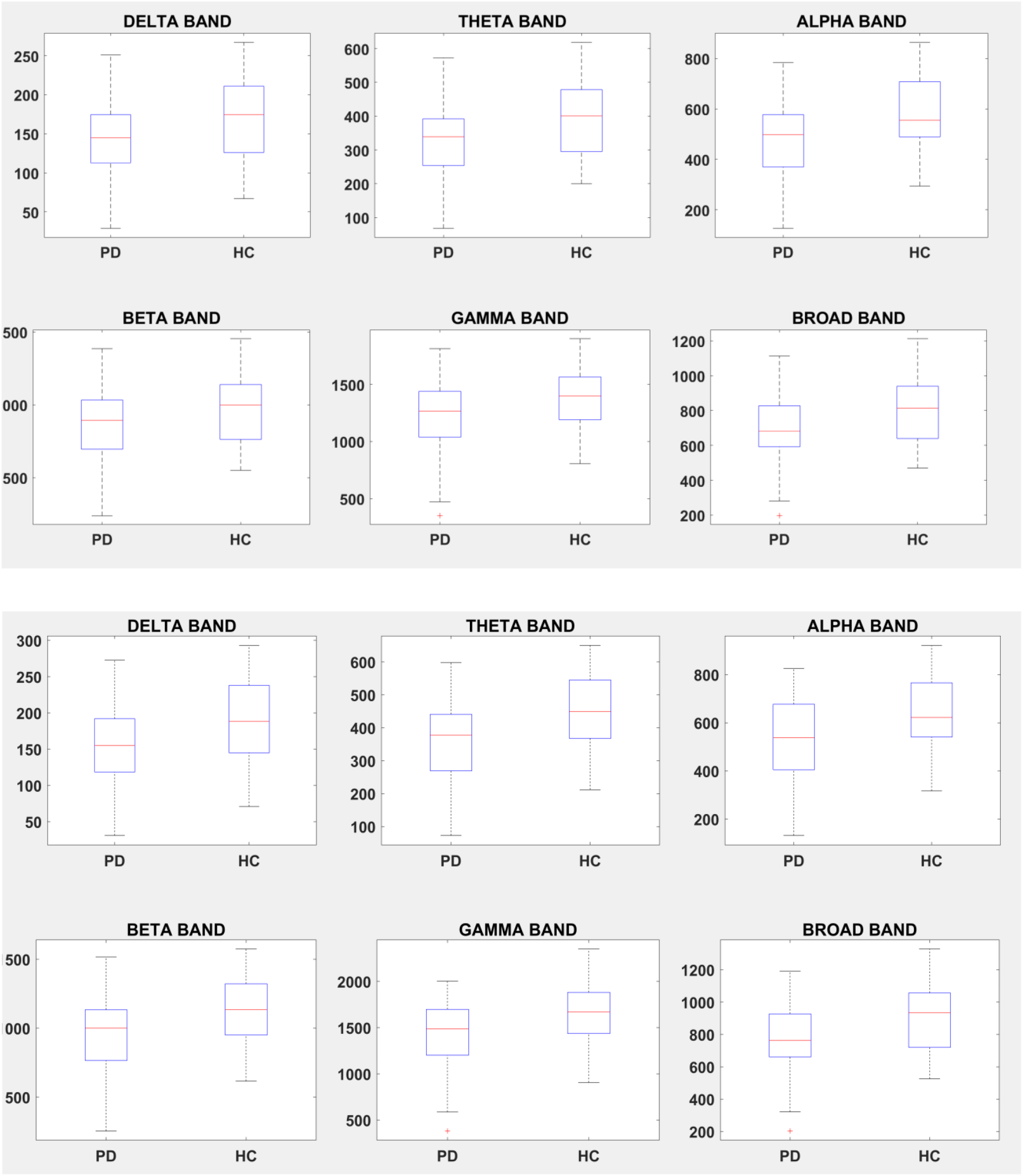
Differences in the size of the functional repertoire in Parkinson patients (PD) and healthy controls (HC), taking into account 90 brain areas (top) and 116 brain areas (bottom).

**Supplementary figure 4.**
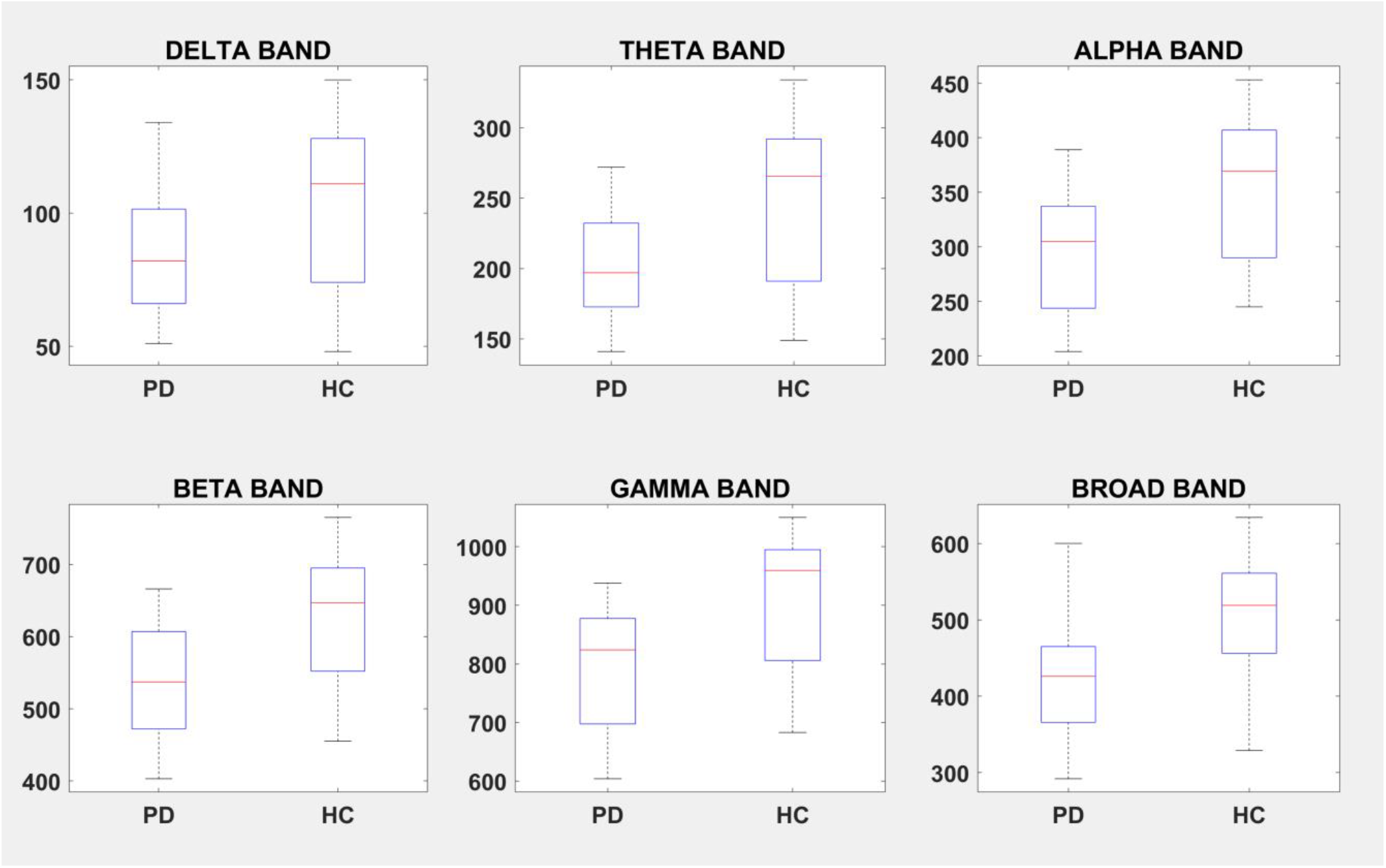
Differences in the size of the functional repertoire in Parkinson patients (PD) and healthy controls (HC), with the same length of data for each participant.

